# cGMP-dependent pathway and a GPCR kinase are required for photoresponse in the nematode *Pristionchus pacificus*

**DOI:** 10.1101/2024.05.28.596172

**Authors:** Kenichi Nakayama, Hirokuni Hiraga, Aya Manabe, Takahiro Chihara, Misako Okumura

## Abstract

Light sensing is a critical function in most organisms and is mediated by photoreceptor proteins and phototransduction. Although most nematodes lack eyes, some species exhibit phototaxis. In the nematode *Caenorhabditis elegans*, the unique photoreceptor protein *Cel*-LITE-1, its downstream G proteins, and cyclic GMP (cGMP)-dependent pathways are required for phototransduction. However, the mechanism of light-sensing in other nematodes remains unknown. To address this question, we used the nematode *Pristionchus pacificus*, which was established as a satellite model organism for comparison with *C. elegans*. Similar to *C. elegans*, illumination with short-wavelength light induces avoidance behavior in *P. pacificus*. Opsin, cryptochrome/photolyase, and *lite-1* were not detected in the *P. pacificus* genome using orthology and domain prediction-based analyses. To identify the genes related to phototransduction in *P. pacificus*, we conducted forward genetic screening for light-avoidance behavior and isolated four light-unresponsive mutants. Whole-genome sequencing and genetic mapping revealed that the cGMP-dependent pathway and *Ppa-grk-2*, which encodes a G protein-coupled receptor kinase (GRK) are required for light avoidance. Although the cGMP-dependent pathway is conserved in *C. elegans* phototransduction, GRK is not necessary for light avoidance in *C. elegans*. This suggests similarities and differences in light-sensing mechanisms between the two species. Using a reverse genetics approach, we showed that GABA and glutamate were involved in light avoidance. Through reporter analysis and suppression of synapse transmission, we identified candidate photosensory neurons. These findings advance our understanding of the diversity of phototransduction in nematodes even in the absence of eyes.

**Author summary:** Nematodes are a highly diverse group of animals found in a wide variety of habitats and sensory systems. In particular, light-induced behavior has been found to differ among species. The photoreceptor and its downstream pathways in *Caenorhabditis elegans* have been identified, revealing unique and distinct characteristics compared to those in other animals. However, the mechanisms of photoreception in other nematodes remain largely unknown. This study focused on the analysis of the photoreception mechanisms in *Pristionchus pacificus*, a species for which many genetic and molecular tools are available. Similar to *C. elegans*, *P. pacificus* also exhibits light avoidance behavior towards short-wavelength light; however, known animal photoreceptor genes could not be identified in the *P. pacificus* genome using bioinformatic approaches. Using forward and reverse genetic approaches, we found that certain genes and neurons are required for light avoidance, some of which are conserved in *C. elegans* photoreception. These results suggest that the light-sensing mechanisms of *C. elegans* and *P. pacificus* are similar, yet there are differences between the two species. These findings highlight the various light-sensing mechanisms in nematodes.

## Introduction

Light sensing is important for many animals that use visual information to avoid predators or unfavorable environments, and to find food sources or mating partners. Most animals, including Cnidaria, Ctenophora, and Bilateria, utilize opsins, which belong to the G protein-coupled receptor (GPCR) superfamily, as photoreceptors and downstream signaling pathways. For example, in vertebrate rods and cones, light is absorbed by the retinal chromophore, which binds to opsins. Isomerization of the retinal chromophore causes a conformational change in opsins, which activates downstream signaling pathways such as G proteins and phosphodiesterases (PDEs). This results in a decrease in cyclic GMP (cGMP) levels and the closure of cyclic nucleotide-gated (CNG) channels. Opsins and their downstream signaling pathways have been extensively studied in a wide range of animals, including aspects such as protein structure, signaling mechanisms, and evolution [1,2]. However, opsin-independent phototransduction mechanisms are limited in the animal kingdom, and the details of these mechanisms remain unclear.

Some nematodes have been observed to exhibit photoresponses despite the absence of eyes [3–6]. In particular, the nematode *Caenorhabditis elegans* displays various responses to short-wavelength light, including avoidance behavior [7,8], stopping pharyngeal pumping [9], and spitting out food [10,11]. Furthermore, *C. elegans* can discriminate between colors [12]. Forward genetic screening using light-avoidance behavior has identified a novel photoreceptor protein, *Cel*-LITE-1 [13,14]. In silico prediction of the protein structure and in vivo ectopic expression analysis suggest that *Cel*-LITE-1 is a member of the 7-transmembrane-domain ion channels (7TMICs) and forms a tetramer [15,16]. Similar to GPCRs, 7TMICs have seven transmembrane domains, but their membrane topology is opposite to that of GPCRs, with the N- and C-termini located intracellularly and extracellularly, respectively [16–20]. Putative binding sites for the chromophore and the aromatic amino acids necessary for light absorption have been identified [15,18]. Although *Cel*-LITE-1 is not predicted to be a GPCR, downstream phototransduction of *Cel*-LITE-1 in *C. elegans* ASJ neurons, one of the photosensory cells, requires G proteins and the cGMP-dependent pathway [7,14]. It is predicted that *Cel-*LITE-1 transduces light stimuli mediated by G-protein α-subunits (*Cel-*GOA-1 and *Cel-*GPA-3) and guanylate cyclases (*Cel-*DAF-11 and *Cel*-ODR-1), resulting in the production of cGMP. Elevated cGMP levels lead to the opening of CNG channels (*Cel*-TAX-2 and *Cel*-TAX-4), causing an influx of calcium ions into the cell. In contrast to vertebrate rods and cones, PDEs (*Cel*-PDE-1, *Cel*-PDE-2, and *Cel*-PDE-5) are not necessary for light response in *C. elegans* [14], suggesting a unique opsin-independent mechanism of phototransduction. However, the mechanism of the light response in other nematode species that lack conventional opsins and LITE-1 remains unknown.

The diplogastrid nematode *Pristionchus pacificus* has been established as a satellite model organism for comparison with *C. elegans* [21–23]. Several genetic tools have been developed for *P. pacificus,* such as an annotated genome [24,25], forward and reverse genetics [21,26], and synaptic connectome in pharynx and amphid neurons revealed using electron microscope [27,28], which provide a suitable model to understand its neural response and behavioral evolution [29–38]. Both *C. elegans* and *P. pacificus* have 12 pairs of amphid neurons, and putative amphid neuronal homologs have been identified between these two species [27]. However, it is likely that the functions of amphid neurons differ between the two species, which is supported by the fact that ciliary terminal structures and the expression of amphid neuron-specific genes vary between the two species [27]. It is currently unknown whether *P. pacificus* has the ability to sense and respond to light.

Here, we found that *P. pacificus* avoids short-wavelength light, although we did not find a conventional opsin cryptochrome/photolyase or *lite-1* in the *P. pacificus* genome. Forward genetic screening revealed that the cGMP-dependent pathway and a GPCR kinase (GRK) are necessary for light avoidance. In addition, a reverse genetic approach has shown that the neurotransmitters gamma-aminobutyric acid (GABA) and glutamate play a role in light avoidance. These genes were expressed in five amphid neurons, and the inhibition of neurotransmission in these amphid neurons reduced light avoidance.

## Results

### *P. pacificus* responds to short-wavelength light

To investigate whether *P. pacificus* possesses light-sensing ability, we established a light avoidance assay for *P. pacificus* based on previous studies on *C. elegans* [7,13]. We illuminated the heads of the worms with different wavelengths of light (UV, blue, and green) for five seconds. In *P. pacificus*, exposure of the head of a forward-moving worm to short-wavelength light halted its forward movement and induced backward movement (Supplemental Movie S1). The percentage of avoidance increased as the light intensity increased, which is similar to *C. elegans* (Fig 1A) [7]. We also illuminated the whole body with different wavelengths of light (UV, blue, green, and red) and observed that *P. pacificus* stopped moving forward and began moving backward under UV and blue light illumination (Fig 1B). Avoidance behavior was not induced by illumination of the entire body with green or red light. A previous study reported that whole-body illumination induces forward movement in *C. elegans* [13], but in our experiments, *C. elegans* moved backward after exposure to light (Fig 1B). For both head and whole-body irradiation, *P. pacificus* exhibited a higher percentage of light-avoidance behavior than *C. elegans*. (Fig 1A, B). These results show that *P. pacificus* has the ability to detect light and is more sensitive to light than *C. elegans*.

**Fig 1.**
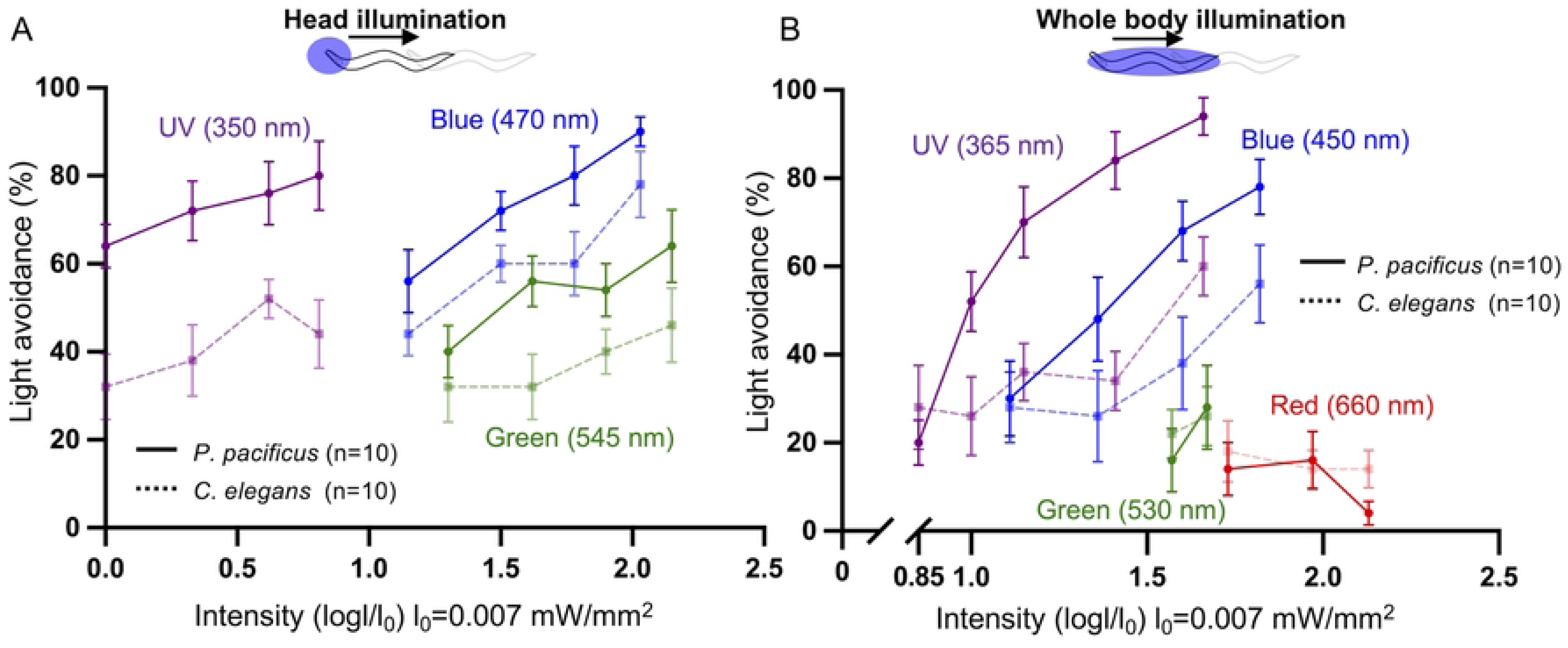
*P. pacificus* exhibited avoidance behavior in response to light illumination. (A, B) Results of light avoidance assays using *C. elegans* (dashed lines) and *P. pacificus* (straight line). UV, blue, or green light illuminated the head (A) or whole body (B) of the worms. *C. elegans* and *P. pacificus* responded to short wavelength light.

### A combination of orthology and domain prediction could not identify putative photoreceptor proteins in *P. pacificus*

We investigated whether *P. pacificus* possesses any known animal photoreceptor proteins by combining orthology and domain prediction, based on previous studies by Pratx et al, 2018 [39] and Brown et al, 2024 [40] (Fig 2A). We obtained the protein sequences of three animal photoreceptor protein families: opsin, cryptochrome/photolyase, and LITE-1 [18,41]. These sequences and their functional domains were used to search for orthologs in *P. pacificus*. Specifically, we evaluated whether each protein in *P. pacificu*s: (1) was orthologous to known photoreceptor proteins, and (2) contained protein domains typical of the respective photoreceptor protein families. To conduct these evaluations, we obtained the protein sequences for 6,040 opsins and 2,249 cryptochromes/photolyases from two previously published reference datasets [42,43]. Additionally, 88 sequences representing the LITE-1 family across 28 species were retrieved from WormBase ParaSite [44] using BLASTP with the *Cel-*LITE-1 sequence as the query. This dataset was expanded by adding *Cel-*EGL-47 (also named *gur-1*), a paralog of *Cel-*LITE-1 (*gur-2*), and the sequence of *Gr28b* in *Drosophila melanogaster*, the closest homolog of *Cel-*LITE-1. *Gr28b* has been implicated in UV light sensing in larvae [45]. The final dataset for the LITE-1 family comprised 95 sequences (S1 Data).

**Fig 2.**
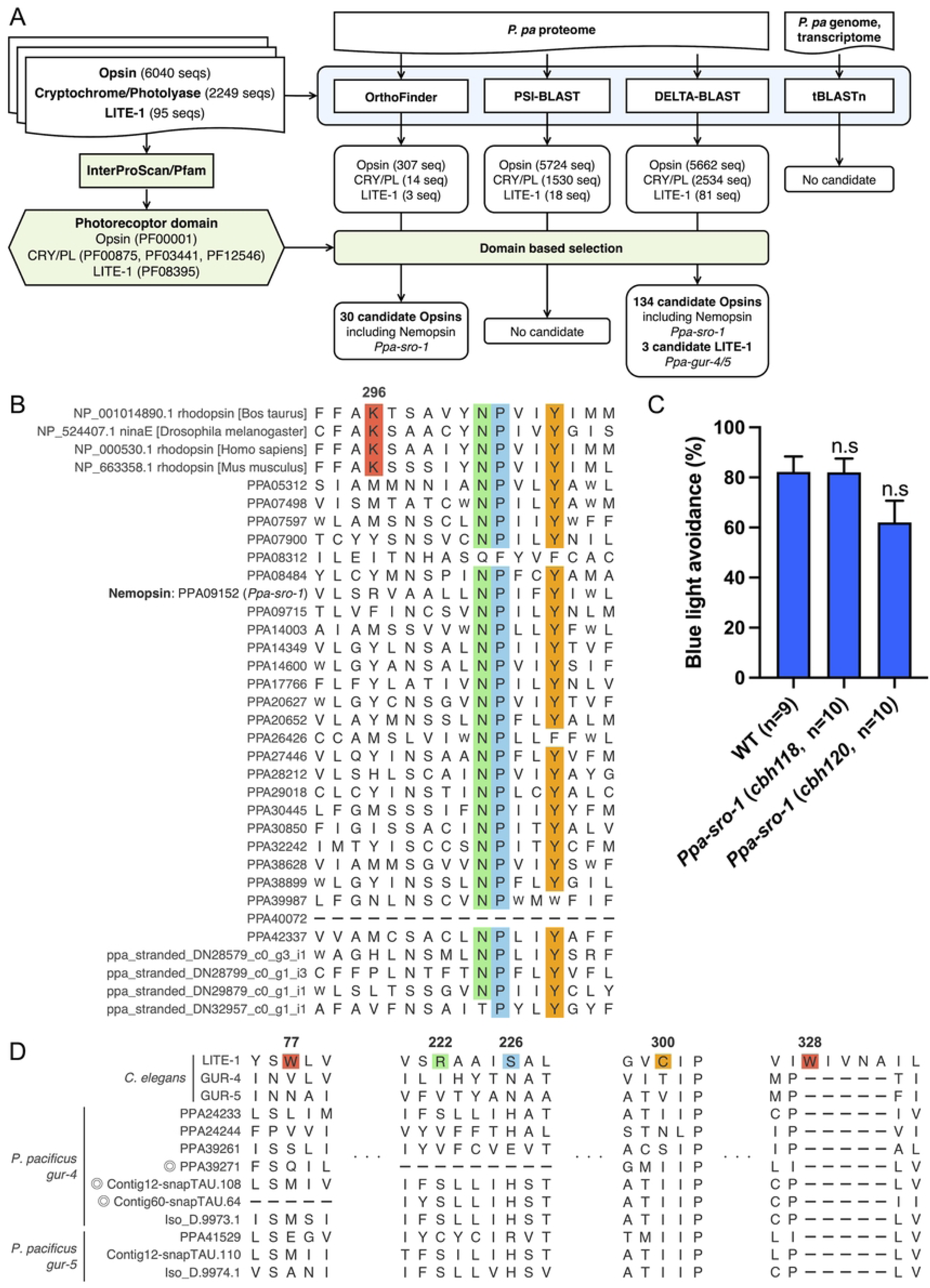
Conserved photoreceptor proteins were not identified in *P. pacificus* using a combination of orthology and domain prediction analyses. (A) Search pipeline for known photoreceptors within the *P. pacificus* multiome. Using InterProScan/Pfam, protein domains were identified in *P. pacificus* proteomes and three photoreceptor families in other animals (Green). CRY/PL; Cryptochrome/Photolyase. Clustering of each photoreceptor family and all *P. pacificus* proteins into orthogroups was performed using OrthoFinder (Blue). PSI-/DELTA-BLAST and tBLASTn were performed to search for known photoreceptors in the *P. pacificus* multiome (Blue). Finally, data outputs from OrthoFinder and BLAST were utilized to identify *P. pacificus* proteins that possess Pfam domains associated with photoreceptors. (B) Sequence alignment of candidate photoreceptor proteins identified by OrthoFinder to the opsin family. While the retinal-binding lysine K296 (*Bos taurus*, red box) is highly conserved among opsins, the candidate GPCRs in *P. pacificus* do not have the conserved lysine. The NPxxY motif, which is highly conserved among GPCRs in their seventh transmembrane domain, is colored with green, blue and orange. (C) Blue light avoidance assay for *Ppa-sro-1* mutants. These mutants showed normal percentage of light avoidance. One-way ANOVA, Dunnett’s multiple comparison tests, compared with wild type. n.s. = not significant.(D) Sequence alignment of *Cel-*LITE-1/GUR-4/5 and *Ppa*-GUR-4/5 in the LITE-1 family. Critical amino acid residues (W77, R222, S226, C300, W328) for photoreception in *Cel-*LITE-1 are not conserved in *Ppa*-GUR-4/5. Circles indicate candidate genes detected in the pipeline.

We identified the functional protein domains of each photoreceptor family using InterProScan domain prediction and the Pfam database (Fig 2A). Each photoreceptor family had a common functional protein domain: opsins were characterized by the 7-transmembrane receptor (rhodopsin family) (PF00001); cryptochrome/photolyase were identified by three domains - DNA photolyase (PF00875), FAD binding domain of DNA photolyase (PF03441), and Blue/Ultraviolet sensing protein C terminal (PF12546); LITE-1 was defined by the 7tm Chemosensory receptor (PF08395).

We performed a BLASTP search using OrthoFinder, targeting all protein sequences of *P. pacificus*, and identified 324 orthogroups. This search yielded 307 hits for opsin, 14 for cryptochrome/photolyase, and three for LITE-1 (S1 Table). Additionally, we conducted both PSI- and DELTA-BLAST searches [46–48] against the *P. pacificus* proteome for each known photoreceptor protein because it is possible that *P. pacificus* photoreceptor proteins are only distantly related to previously known photoreceptor proteins and might not be detectable by BLASTP alone. PSI-BLAST identified 5724 hits for opsin, 1530 for cryptochrome/photolyase, and 18 for LITE-1 (S2 Table). DELTA-BLAST, which uses conserved protein domain databases, detected 5662 hits for opsin, 2534 for cryptochrome/photolyase, and 81 for LITE-1 (S2 Table). Furthermore, to determine whether there were any indications of the loss of known photoreceptor protein genes within the genome, we used tBLASTn to search the photoreceptor protein reference dataset against both the genome (El Paco Assembly) [25] and the transcriptome (El Paco V3) of *P. pacificus*. However, the majority of tBLASTn hits overlapped with those identified by PSI-BLAST, and many of the genomic region hits corresponded to either introns or non-coding sequences (S2 Table).

We then checked whether these candidate genes encode the protein domains of characteristic of each photoreceptor. Among them, *P. pacificus* proteome has 30 genes in OrthoFinder and 134 genes in DELTA-BLAST, respectively, with conserved domains characteristic of opsin family specific domains (PF00001), no genes with the cryptochrome/photolyase, and 3 genes with *lite-1* families (Fig 2A). However, the highly conserved lysine residue among opsins (K296, as found in *Bos taurus*), which enables opsins to bind retinal and is crucial for photoreception [49], was not conserved in these protein sequences (Fig 2B, we showed only 30 sequences identified in OrthoFinder). Among these genes, we identified *Ppa-sro-1*, which belongs to the nemopsin family. Nemopsins are a family of chromopsins that includes peropsins, RGR-opsins, and retinochromes, some of which function as photoreceptor proteins; however, nemopsins feature a substitution of conserved lysine to arginine [42] (Fig 2B). In *C. elegans*, *Cel-sro-1* is expressed in ADL chemosensory neurons and SIA motor neurons [50], but the functions of *Cel-sro-1* and *Ppa-sro-1* are unknown. The percentage of light avoidance behavior in the *Ppa-sro-1* mutant was similar to that in the wild type (Fig 2C), suggesting that *Ppa-sro-1* does not have photoreceptor functions.

DELTA-BLAST using LITE-1 as a query detected 81 genes (S3 Table), three of which were annotated using protein domain prediction by InterProScan (Fig 2A). To assess their potential as photoreceptors, we compared the crucial amino acid residues for photoreception in *Cel-*LITE-1 (W77, R222, S226, C300, and W328) [15,18] against these three genes and other *Ppa-gur-4/5* homologs because all three genes are homologous to *Cel-gur-4/5*. However, we found that none of the *Ppa-gur-4/5* genes encode for the amino acid residues essential for photoreception (Fig 2D). These results suggest that, despite using a combined approach of orthology clustering, extensive BLAST searches, domain prediction, and sequence alignment to compare conserved residues, conserved photoreceptor proteins in *P. pacificus* could not be identified.

### cGMP-dependent pathway is required for light avoidance in *P. pacificus*

To identify the genes responsible for light avoidance in *P. pacificus*, we conducted forward genetic screening using light avoidance behavior. We mutagenized wild-type *P. pacificus* using ethyl methanesulfonate (EMS) and tested the light-avoidance behaviors of F2 and F3 animals.

We screened more than 20,000 strains and identified four light-unresponsive mutants. These mutants exhibited a decreased percentage of light avoidance (Fig 3A). Whole-genome sequencing revealed that *cbh37*, *cbh44*, and *cbh87* harbored mutations in *Ppa-daf-11*, which encodes guanylate cyclase (Fig 3B). This gene is a one-to-one ortholog of *Cel-daf-11* and is involved in the phototransduction of the ASJ sensory neuron in *C. elegans* (Fig 3C). This led us to hypothesize that *P. pacificus* uses a similar phototransduction to *C. elegans*. Since the G-protein α-subunit and the cGMP-dependent pathway are used in *C. elegans* phototransduction [7,14], we generated mutants of the G-protein α-subunit (*Ppa-goa-1*; *Ppa-gpa-3* double), guanylate cyclases (*Ppa-odr- 1*, *Ppa-daf-11,* and *Ppa-odr-11*; *Ppa-daf-11* double), CNG channels (*Ppa-tax-2, Ppa-tax-4*, and *Ppa-tax-2*; *Ppa-tax-4* double), and cGMP specific phosphodiesterase (*Ppa-pde-1*; *Ppa-pde-2*; *Ppa-pde-3*; *Ppa-pde-5* quattro) using the CRISPR/Cas9 system (Fig 3C). While G-protein α- subunit mutants displayed normal light avoidance, mutants of guanylate cyclases, CNG channels, and the phosphodiesterases (PDEs) decreased the percentage of light avoidance (Fig 3D). These results support our hypothesis that *P. pacificus* and *C. elegans* exhibit conserved cGMP-dependent phototransduction.

**Fig 3.**
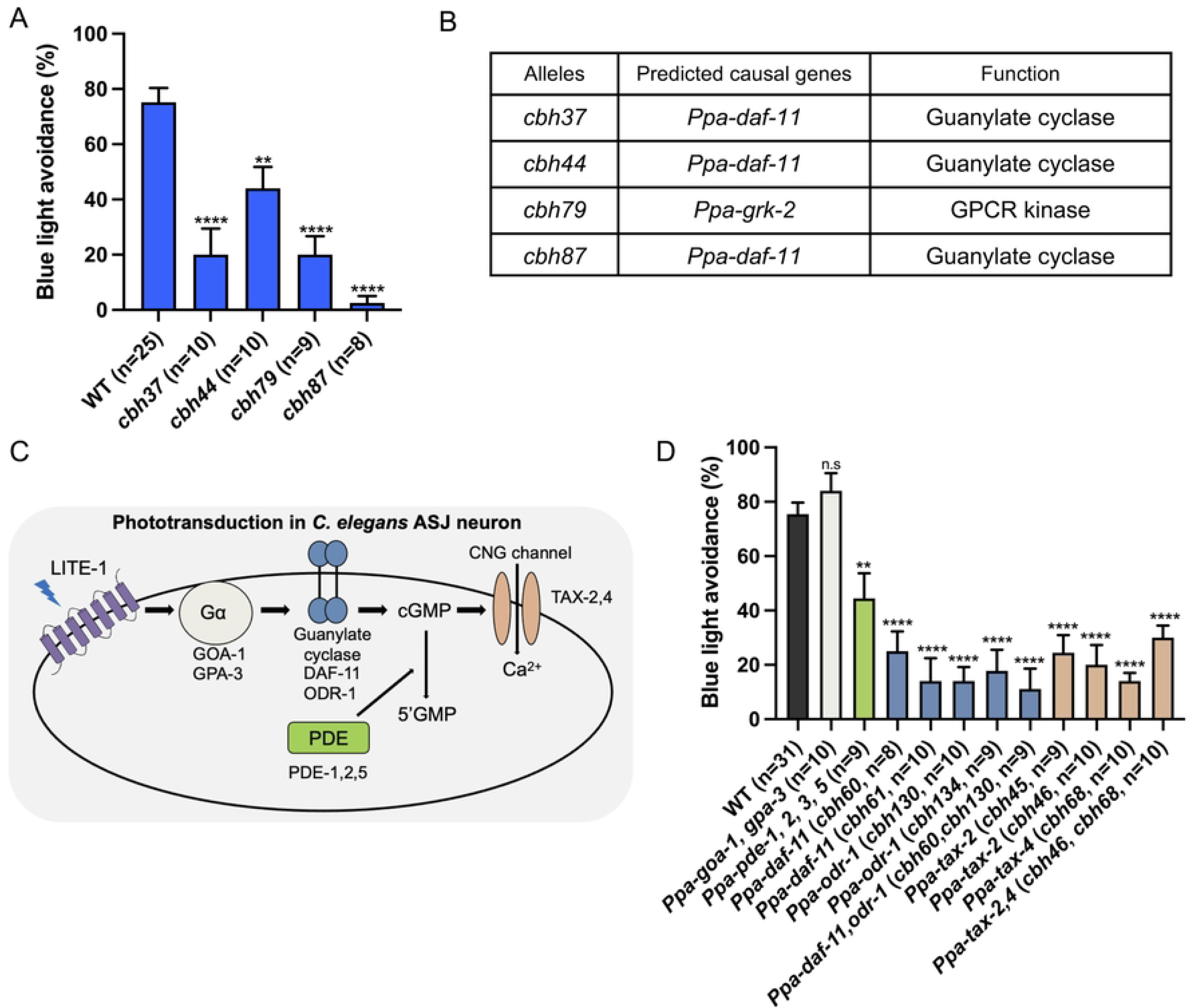
Light avoidance requires the cGMP-dependent pathway in *P. pacificus.* (A) Blue light avoidance assay for light unresponsive mutants that were isolated by forward genetic screening. These mutants had decreased light-induced avoidance response. (B) List of predicted causal genes. (C) Scheme of phototransduction in the *C. elegans* ASJ neuron. (D) Blue light avoidance assay for mutants of cGMP-dependent pathway genes. Mutants of guanylate cyclases, CNG channels, and PDEs had defect in light avoidance in *P. pacificus*. G-protein α- subunit mutants displayed normal light avoidance. One-way ANOVA, Dunnett’s multiple comparison tests, compared with wild type. n.s = not significant, \*\**P* < 0.01, \*\*\*\**P* < 0.0001.

### GPCR kinase is required for light avoidance in *P. pacificus* but not in *C. elegans*

Genetic mapping revealed that an EMS mutant, *cbh79*, harbored a mutation in *Ppa-grk-2*, which encodes a G protein-coupled receptor kinase (GRK) (Fig 3B). GRKs play a crucial role in the regulation of GPCRs by phosphorylating activated GPCRs, subsequently leading to the binding of arrestin to GPCR. This interaction induces endocytosis and desensitization [51] (Fig 4A). In vertebrate rod and cone cells, GRKs phosphorylate light-activated opsins, which are crucial for photoreception [52–56]. Both *P. pacificus* and *C. elegans* possess two GRK genes (*grk-1* and *grk- 2*) and an arrestin gene (*arr-1*). In *C. elegans*, *Cel-grk-2* is essential for chemosensation [57,58], but *Cel-grk-1* and *Cel-grk-2* mutants showed normal light avoidance (Fig 4B), which is consistent with the fact that the *C. elegans* photoreceptor *Cel*-LITE-1 is a member of 7TMICs rather than GPCRs [15,16,20].

**Fig 4.**
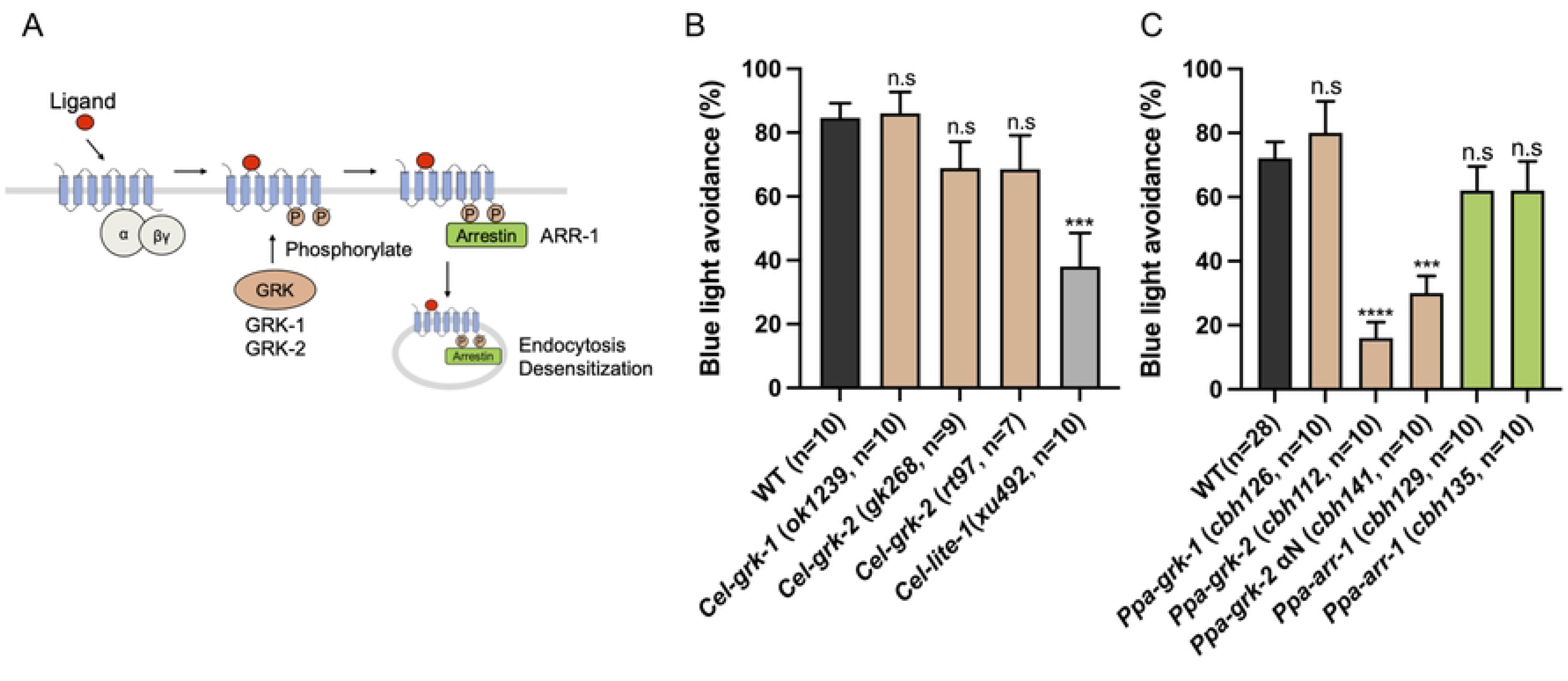
GPCR kinase, *Ppa*-GRK-2 is required for light avoidance. (A) Schematic of GPCR desensitization by GRKs. After a ligand binds to a GPCR, G proteins are released from the activated GPCR to transmit signaling. GRKs phosphorylate the activated GPCR, promoting the binding of arrestin to the receptor. GPCR is endocytosed and desensitized. (B) Blue light avoidance assay for GRK and arrestin mutants in *C. elegans*. These mutants displayed normal light avoidance. The *Cel-lite-1* mutant is used as a negative control. (C) Blue light avoidance assay for GRK and arrestin mutants in *P. pacificus*. *Ppa-grk-2* mutants had decreased light avoidance. One-way ANOVA, Dunnett’s multiple comparison tests, compared with wild type. n.s = not significant, \*\*\**P* < 0.001, \*\*\*\**P* < 0.0001.

We generated mutants in *Ppa-grk-1*, *Ppa-grk-2*, and *Ppa-arr-1* using the CRISPR/Cas9 system in *P. pacificus*. We found that the *Ppa-grk-2* mutants, but not the *Ppa-grk-1* or *Ppa-arr-1* mutants, exhibited decreased light avoidance (Fig 4C), suggesting different roles for *Ppa*-GRK- 1 and *Ppa*-GRK-2. We also generated a *Ppa-grk-2* mutant (*cbh141*) lacking a part of the αN domain that stabilizes its binding to activated GPCRs. The absence of the αN domain resulted in a reduction of light avoidance (Fig 4C), implying that the potential involvement of GPCR phosphorylation in light avoidance mechanism in *P. pacificus*. Together, these results indicate that the GRK-2, particularly its αN domain, is necessary for light avoidance in *P. pacificus* but not in *C. elegans*.

### Neurotransmitters GABA and glutamate are required for light avoidance

To identify the genes involved in light avoidance, we focused on neurotransmitters. In *C. elegans*, neurotransmitters, glutamate, and glutamate receptors are involved in light-avoidance behavior [8]. We performed a blue light avoidance assay with head illumination using mutants of the following neurotransmitter-related genes: *Ppa-tph-1* encoding a tryptophan hydroxylase responsible for serotonin synthesis, *Ppa-tdc-1* encoding a tyrosine decarboxylase required for tyramine and octopamine synthesis, *Ppa-cat-2* encoding a tyrosine hydroxylase required for dopamine synthesis, *Ppa-unc-25* encoding a GABA synthesis enzyme; and *Ppa-eat-4* encoding a vesicular glutamate transporter. The mutants *Ppa-tph-1*, *Ppa-tdc-1*, and *Ppa-cat-2* did not show a significant difference in the percentage of light avoidance compared to the wild-type (Fig 5). In contrast, mutants of *Ppa-unc-25* and *Ppa-eat-4* exhibited a decrease in the percentage of light avoidance (Fig 5). These findings indicated that both GABA and glutamate are involved in light avoidance in *P. pacificus*.

**Fig 5.**
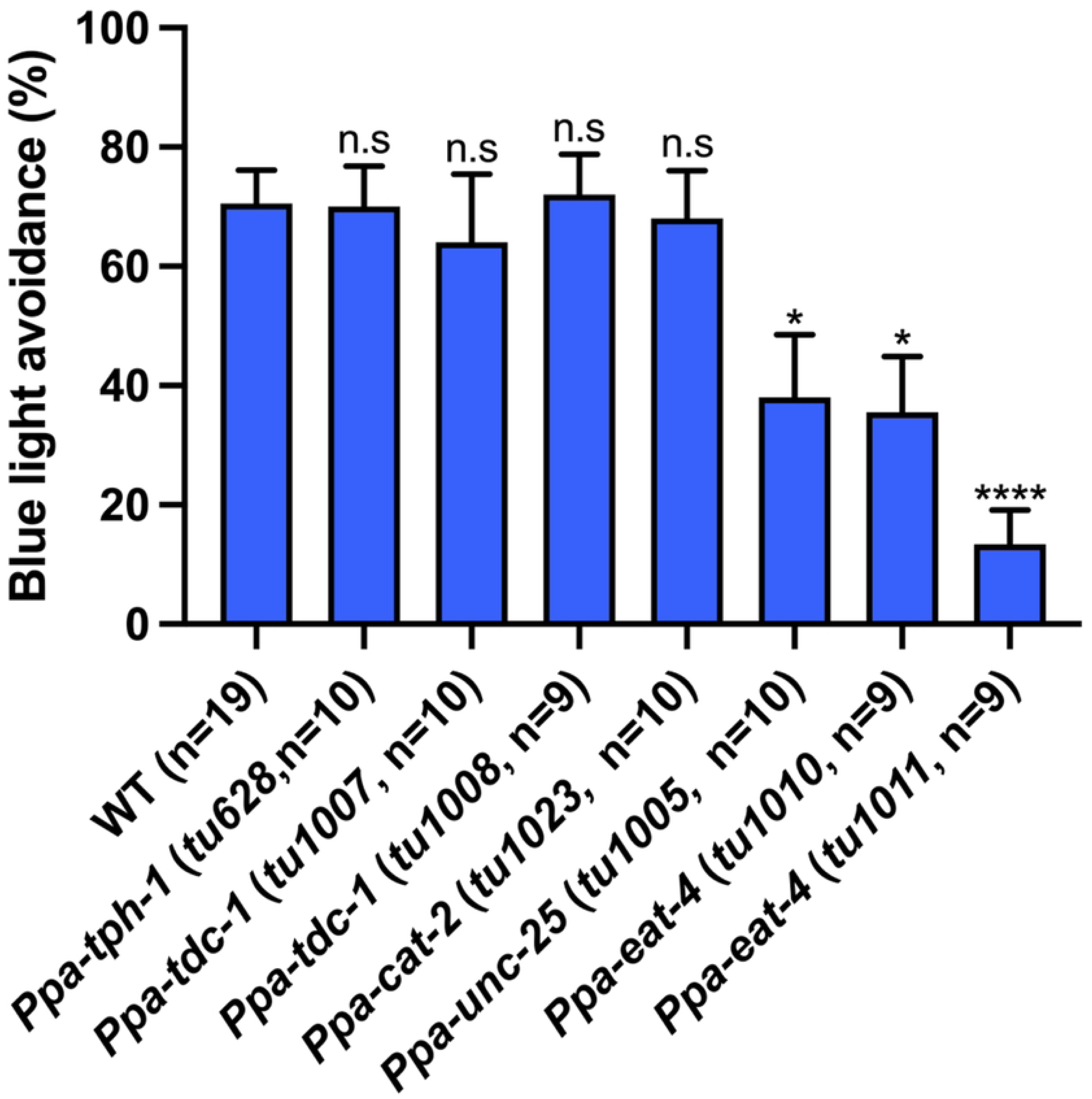
GABA and glutamate are required for light avoidance. Blue light avoidance assay for neurotransmitter-related mutants. *Ppa-unc-25* and *Ppa-eat-4* mutants exhibit decreased light avoidance. One-way ANOVA Dunnett’s multiple comparison tests, compared with wild type. n.s = not significant, \**P* < 0.05, \*\*\*\**P* < 0.0001.

### Amphid neurons expressing *Ppa-daf-11, tax-2, tax-4,* and *grk-2* mediate light avoidance behavior

To identify the photosensory neurons, we generated transgenic reporter lines for *Ppa-grk-2*, *Ppa-tax-2*, and *Ppa-tax-4*. We found that *Ppa-tax-2* and *Ppa-tax-4* were expressed in the head and amphid neurons (Figs 6A and 6B). *Ppa-grk-2* showed a broader expression pattern, including in amphid neurons, pharyngeal muscles, head neurons, body wall muscles, and ventral nerve codes (Fig 6C and S1A, B Fig). Since in *C. elegans*, amphid neurons such as ASJ, ASK, and ASH neurons are photosensory neurons, we focused on the amphid neurons of *P. pacificus* for further cell identification. Previous studies have shown that *Ppa-daf-11* is expressed in AM1, 3, 4, 5, and 8 neurons [59]. Our reporter lines revealed that *Ppa-tax-2* and *Ppa-tax-4* were expressed in AM1, 3, 4, 5, 6, 7, 8, and 12 neurons, while *Ppa-grk-2* was expressed in AM1, 2, 3, 4, 5, 6, 8, 9, 10, and 11 neurons (Fig 6D). AM1, 3, 4, 5, and 8 neurons expressed all the four genes (Fig 6D). These results suggest that these amphid neurons are potential photosensory neurons.

**Fig 6.**
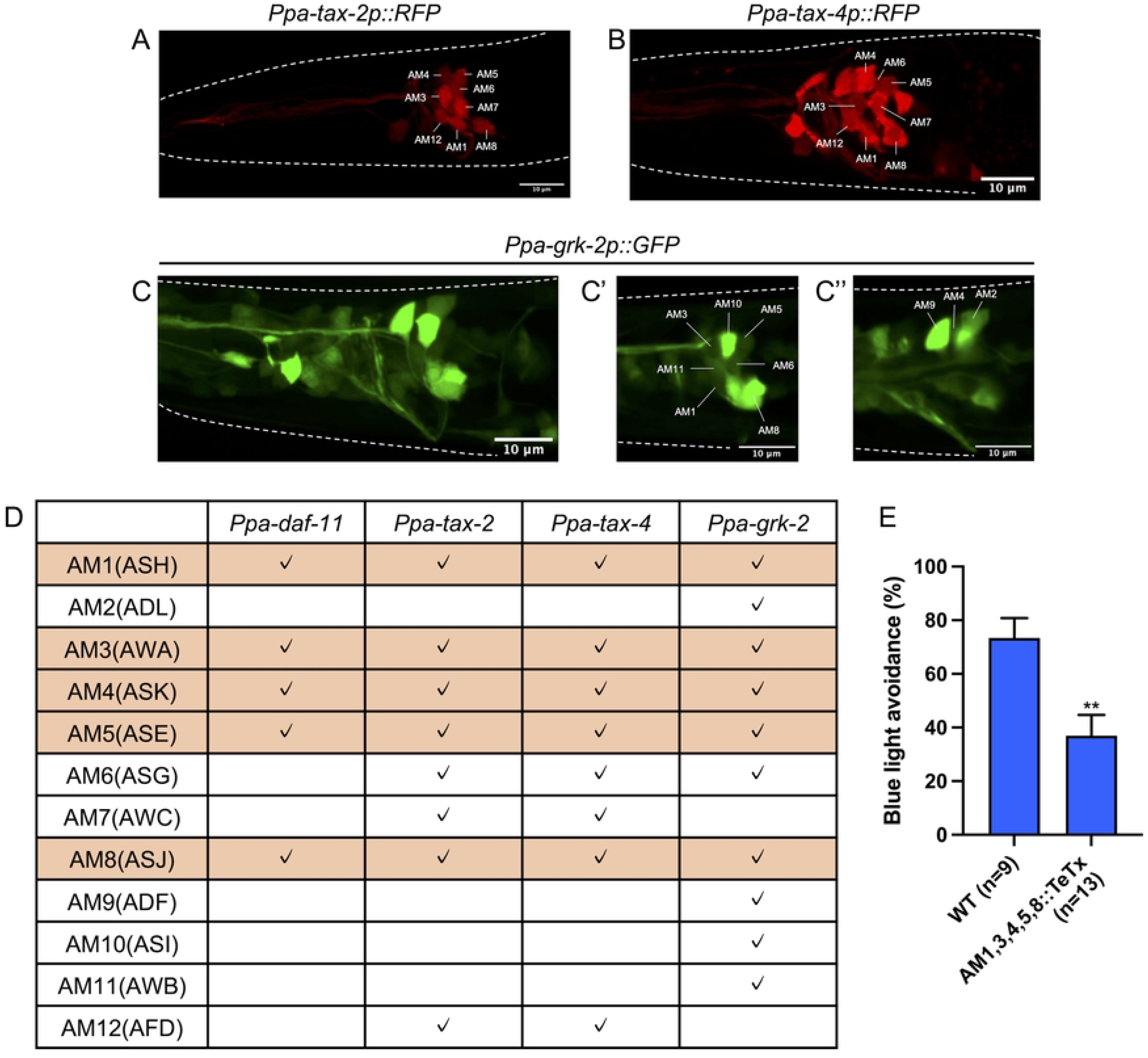
Amphid neurons expressing guanylate cyclase, CNG channels, and GRK-2 are important for light avoidance. (A–C) Representative fluorescence images of *Ppa-tax-2p::RFP* (A), *Ppa-tax-4p::RFP* (B), and *Ppa-grk-2p::GFP* (C) in the head region. Anterior is left and dorsal is up. Scale bars = 10 μm. (A–C) are maximum projection images. (C′) and (C′′) are single focal plane images. (D) Summary of the expression pattern of phototransduction genes. Check marks indicate that the reporter fluorescent proteins were expressed in the corresponding cells in more than 80% individuals. All four phototransduction genes we examined were expressed in AM1, 3, 4, 5, and 8 neurons (orange columns). (E) Blue light avoidance assay for worms expressing tetanus toxin in AM1, 3, 4, 5, and 8 neurons using the *Ppa-daf-11* promotor. The transgenic animals had decreased light avoidance. Student’s *t*-test. \*\**P* < 0.01.

Amphid neurons in *P. pacificus* are ciliated [27] and are important for detecting environmental stimuli [38,60,61]. To investigate the role of cilia in light detection, we examined light avoidance in cilia-related mutants, some of which exhibit abnormal morphogenesis in amphid neurons [38,60]. Mutants of *Ppa-daf-19,* encoding a regulatory factor X transcriptional factor, and several intraflagellar transport components including IFT-B (*Ppa-osm-1*), BBsome (*Ppa-osm-12*), Kinesin-2 (*Ppa-klp-20*), and Dynein-2 (*Ppa-che-3*), exhibited normal light avoidance (S2 Fig), showing that cilia are not necessary for light avoidance in *P. pacificus*.

To examine whether these amphid neurons are necessary for light avoidance, we used tetanus toxin to inhibit the release of neurotransmitters and neuropeptides in specific cells [62]. We expressed the codon-optimized tetanus toxin in AM1, 3, 4, 5, and 8 neurons by utilizing the *Ppa-daf-11* promoter. This transgenic strain exhibited reduced light-avoidance (Fig 6E). Taken together, these results suggest that these cells are candidate photoreceptors that mediate light avoidance in *P. pacificus*.

## Discussion

*C. elegans* utilizes a unique *Cel-*LITE-1-dependent photoreception instead of conventional animal photoreceptor proteins such as opsin and cryptochrome/photolyase. However, it remains unclear how other nematodes sense light. In the present study, we identified the genes and neurons that mediate light-avoidance behavior in the diplogastrid nematode *P. pacificus*. Although the combination of orthology and domain prediction could not identify opsin, cryptochrome/photolyase, or *lite-1* in its genome (Fig 2), *P. pacificus* responds to light, suggesting that *P. pacificus* possesses photoreceptor proteins that differ from known photoreceptor proteins. Using forward and reverse genetic approaches, we found that the cGMP-dependent pathway, GRK-2, GABA, glutamate, and some amphid neurons mediate light-avoidance behavior in *P. pacificus*. Because glutamate, the cGMP-dependent pathway, and amphid neurons, but not GRKs, are also used for photoresponses in *C. elegans* [7,8,14] (Fig 4B), *C. elegans* and *P. pacificus* have similar but different light-response mechanisms (Fig 7).

**Fig 7.**
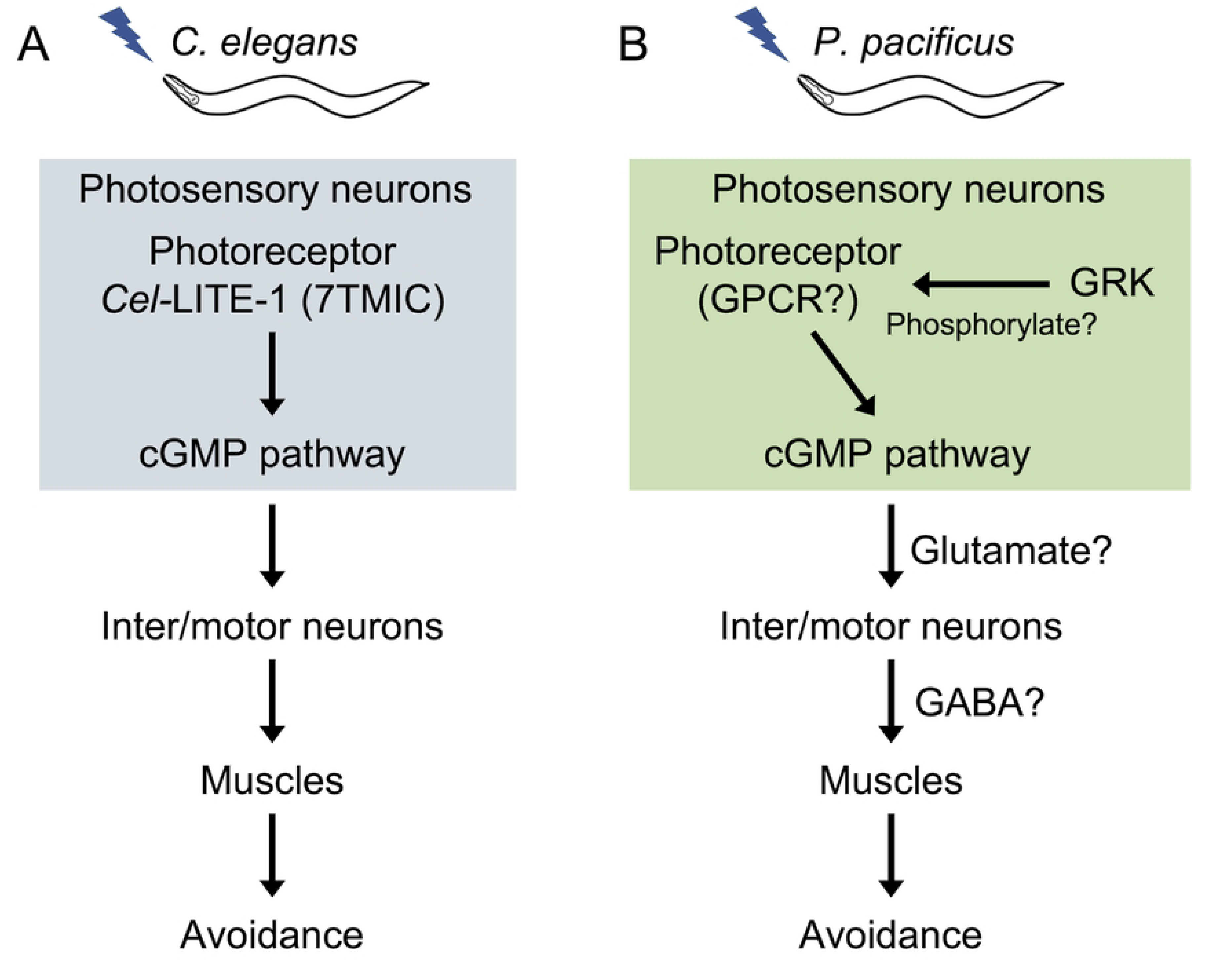
Proposed regulatory models of light avoidance behavior in *C. elegans* and *P. pacificus.* (A) In *C. elegans*, *Cel*-LITE-1, a photoreceptor protein in photosensory neurons (ASJ and ASK) receives light and transmits the signal to the downstream cGMP-dependent pathway. Signals from the photosensory neurons are transmitted to the interneurons and motor neurons. (B) In *P. pacificus*, unknown photoreceptor proteins in the photosensory neurons (AM1, 3, 4, 5, or 8) detect light and transmit the signal to the downstream cGMP-dependent pathway. Signals from photosensory neurons are transmitted to interneurons and motor neurons via glutamate or GABA, leading to the muscle contraction and avoidance behavior. GRKs may phosphorylate photoreceptor proteins.

The phototransduction of *P. pacificus* is similar in some aspects to the previously known phototransduction in *C. elegans*. Although *Cel*-LITE-1 has been proposed to function as a light- activated ion channel [15], the G-protein α-subunit and the cGMP-dependent pathway is required for phototransduction downstream of *Cel*-LITE-1 in the ASJ and ASK neurons [7,14]. In *C. elegans*, phototransduction does not require PDEs in ASJ neurons [14]. In *P. pacificus*, the guanylate cyclases (*Ppa-odr-1, Ppa-daf-11*) and CNG channels (*Ppa-tax-2, 4*) are required for light avoidance behavior (Fig 3D), and mutants of *Ppa-pde-1, 2, 3*, and *5* showed a reduced percentage of light avoidance (Fig 3D). These results suggest that, although they share the same cGMP signaling pathway, there may be differences in the activation mechanism of the cGMP signaling pathway.

The cGMP-dependent pathway is crucial for opsin-dependent phototransduction in various animals, including vertebrate rods and cones, marine mollusks, scallops, and lizard parietal eyes [41]. In vertebrate rod and cone cells, GPCR opsins are phosphorylated and desensitized by GRKs [52–56]. The percentage of light avoidance decreased in the *Ppa-grk-2* knock-out mutants and mutants lacking a part of the αN domain, which is thought to be important for GPCR binding (Fig 4C). These results suggest that the photoreceptor in *P. pacificus* is a GPCR that is phosphorylated by *Ppa*-GRK-2. Nemopsin, encoded by *Ppa-sro-1*, is a recently described opsin that lacks a conserved lysine [42]. Nemopsin has the closest amino acid sequence to opsin among all the GPCRs of *P. pacificus*. However, our mutant analysis revealed that *Ppa-sro-1* mutants exhibited normal light avoidance (Fig 2C). It is possible that nemopsin functions redundantly with other photoreceptor proteins, but it is likely that at least one protein that differ from known photoreceptors functions as a photoreceptor in *P. pacificus*. Notably, the amino acid sequence of LITE-1, which belongs to the 7TMIC superfamily [15,16,20], differs greatly from that of existing photoreceptor proteins. Recently, 7TMICs, which were thought to be conserved only in invertebrates, were revealed to be ancient, conserved proteins with highly divergent amino acid sequences by the identification of homologous proteins based on protein structure [16,20]. Future structure-based analyses may allow the identification of photoreceptor proteins that cannot be identified by amino acid sequence homology.

We identified candidates for the five photosensory neurons, AM1, 3, 4, 5 and 8. These neurons correspond to the ASH, AWA, ASK, ASE, and ASJ neurons in *C. elegans* [27]. The ASK and ASJ neurons are photosensory neurons that utilize the cGMP-dependent pathway for phototransduction [7,14]. Therefore, the corresponding AM4 and 8 neurons are potential candidates as photosensory neurons. In the future, photosensory neurons could be identified through calcium imaging or electrophysiological recordings.

We discovered that both *Ppa-unc-25*, a synthetic enzyme of the neurotransmitter GABA, and *Ppa-eat-4*, a vesicular glutamate transporter, are required for light avoidance in *P. pacificus* (Fig 5). In *C. elegans*, *Cel-unc-25* mutants display hypercontraction of body muscles, excessive flexion of the head while foraging, and a significant decrease in contraction of enteric muscles [63]. In addition, the body bending rate was reduced in the *Ppa-unc-25* in *P. pacificus* (Onodera et al, unpublished). Based on these results, it is likely that the *Ppa-unc-25* mutant is defective in avoidance behavior rather than in the regulation of photoreception. Glutamate and glutamate receptors are required for UV light avoidance behavior in *C. elegans* [8]. The photosensory neurons of *C. elegans*, specifically ASK and ASH neurons, are glutamatergic [64]. Additionally, the photoreceptor cells in the mammalian retina are glutamatergic [65]. Although it is not known which cells are glutamatergic neurons in *P. pacificus*, it is possible that AM1, 3, 4, 5, and 8 neurons are glutamatergic and that glutamate is released from these cells to transmit the light stimuli downstream.

Orthology and domain analyses did not identify a photoreceptor protein in *P. pacificus* (Fig 2). This suggested that *P. pacificus* contains a novel photoreceptor protein. Furthermore, the zoonotic nematode *Dirofilaria immitis* displays positive phototaxis towards infrared light [4]. Although *Cel-*LITE-1 detects UV and blue light, it does not play a role in the perception of longer light wavelengths [15,18]. Therefore, *D. immitis* might possess novel photoreceptor proteins. Thus, nematodes possess various photosensory mechanisms and can serve as valuable models for studying the evolution of photoreception.

## Methods

### Strains

The strains used in this study are listed in S4 Table. *C. elegans* and *P. pacificus* were maintained at 20 °C on Nematode Growth Medium (NGM) agar plates with *Escherichia coli* OP50 as previously described [22,66].

### Light avoidance assay

The light avoidance assay was conducted as previously described [7] with some modifications. All assays were performed in a blinded manner. One-day adult hermaphrodite worms were placed individually on NGM plates covered with a thin bacterial lawn of freshly seeded OP50 and left in the dark for at least 10 min before the assay. For head illumination, in Fig 1A, a fluorescent stereomicroscope (Leica, 165 FC) was connected to a mercury lamp (Leica, EL6000) and the head of the nematode that moved forward was illuminated using a fluorescent filter and an objective lens (Leica, 10450028). Light intensity was adjusted by manipulating the amount of light emitted from the mercury lamp. The following fluorescent filters and wavelengths were used: UV (350 nm), Leica ET UV LP, 1045609; blue (470 nm), Leica ET GFP, 10447408; green (545 nm), Leica ET DSR, 10447412. For other experiments, the LED light source from a fluorescence stereomicroscope (ZEISS, Discovery V20) was illuminated through a fluorescence filter (ZEISS, filter set 38 HE, 470±20 nm, 0.24 mW/mm^2^). For whole-body illumination, light from an LED source (Optocode LED-EXSA) was delivered to the entire body of each nematode. To use light of different wavelengths, we changed the LED head accordingly (red, EX-660; green, EX-530; blue, EX-450; UV, EX-365). When nematodes ceased forward movement and began backward movement within 5 seconds after light irradiation, it was considered as “light-avoidance behavior.” In a previous *C. elegans* study [7], light-avoidance behavior was defined as backward movement within 3 s of light irradiation. However, in this study, we defined it as within 5 s because the locomotion speed of *P. pacificus* is slower than that of *C. elegans*. In Fig 3C, because only *C. elegans* was assayed, backward movement within 3 s was defined as light-avoidance behavior. For each individual, the light avoidance assay was performed five times at 10 min intervals after each assay. A red filter (Kenko 158371) was used to minimize the impact of white light from the lower part of the microscope.

The light intensity was measured using an optical power meter (HIOKI, 3664) with an optical sensor (HIOKI, 9742) divided by the illuminated area. Except for Fig 1, the intensity of blue light (470 nm) was 0.24 mW/mm^2^ (*P. pacificus*) or 1.83 mW/mm^2^ (*C. elegans*) for the light avoidance assay.

### Protein sequence collection

We used previously published datasets and BLASTP sequence searches to collect a reference dataset for photoreceptor protein families. Specifically, for the opsin and cryptochrome/photolyase families, we utilized the data collected by Gühmann et al, 2022 [42] and Deppisch et al, 2022 [43]. For the LITE-1 family, we performed BLASTP (https://parasite.wormbase.org/Multi/Tools/Blast) searches on all nematode species registered in WormBase ParaSite (Version: WBPS18, https://parasite.wormbase.org) [44] using the *C. elegans* LITE-1 (WBGene00001803) protein sequence as query. From this result, we obtained 88 sequences using BioMart (https://parasite.wormbase.org/biomart/martview/). Furthermore, we obtained and added the sequences of *Cel-*EGL-47 (WBGene00001211), *Cel-*GUR-3 (WBGene00001804), and *D. melanogaster* Gr28b (FBgn0045495, from FlyBase; https://flybase.org) which are registered as paralogs or orthologs of *Cel-*LITE-1 in WormBase (Version: WS291, https://wormbase.org) [67,68] *melanogaster* Gr28b (FBgn0045495, from FlyBase; https://flybase.org) [69] which are registered as paralogs or orthologs of *Cel-*LITE-1 in WormBase (Version: WS291, https://wormbase.org) [67,68], to the LITE-1 family (S1 Data). These three protein sequence files were used as inputs for subsequent analyses.

### Exploration of putative photoreceptor proteins

Protein domain searches were performed on the photoreceptor protein reference dataset and all proteins of *P. pacificus* based on the Pfam database (version 36.0) [70] using InterProScan (version 98.0, option: -dp -appl Pfam) [71]. Protein domains that were functionally important as photoreceptor proteins in the reference dataset were isolated and searched in all protein domain dataset of *P. pacificus*.

Orthology clustering with OrthoFinder (Version: 2.5.5, option: -S blast -M msa) [72] was performed on the three photoreceptor protein reference datasets and the *P. pacificus* proteome (El paco V3, http://pristionchus.org) [73] to obtain orthogroup data (S1 Table). Orthogroups from OrthoFinder and domain prediction data from InterProScan were integrated to search for photoreceptor protein candidates from all proteins of *P. pacificus*.

Given that BLASTP, as used in OrthoFinder, may not detect remotely homologous genes, we performed PSI- and DELTA-BLAST (BLAST+, Version: 2.15.0; option: - comp_based_stats 1) for searching photoreceptor protein families in the proteome of *P. pacificus* (S2 Table). PSI- and DELTA-BLAST searches were iterated three times with the threshold set at 1e-3.

To determine whether any traces of known photoreceptor proteins remained in the genome, we performed tBLASTn (BLAST+, Version: 2.15.0) (option: -evalue 1e-3) on the genome and transcriptome of *P. pacificus*, using each photoreceptor protein reference dataset as a query.

### Sequence alignment

The protein sequences used for sequence alignment were candidate protein sequences from a homology search using OrthoFinder/BLAST and protein domain prediction using InterProScan. Sequences were aligned with MAFFT (version 7.525; [74] using the "--auto" option, specifically employing the L-INS-i method. The aligned sequences were visualized in R (Version 4.3.2) using the ggmsa package (Version: 1.3.4) [75].

### Genetic screen for light-unresponsive mutants

*P. pacificus* PS312 was mutagenized with ethyl methanesulfonate (EMS), as described previously [76]. Two methods were used to screen the light-unresponsive mutants. In the first method, the F1 worms were individually transferred onto *E. coli* plates. When the F2 animals reached the adult stage, a light-avoidance assay was conducted once per individual, with 10 individuals per plate. F2 strains exhibiting light avoidance at a frequency of 40% or less were selected, and F2 worms were individually transferred to *E. coli* plates. After a few days, the F3 worms were again tested for light avoidance. The mutants with impaired locomotion were excluded. In the second method, plates containing P0 were left for several days and F2 or F3 individuals were transferred individually to *E. coli* plates. Subsequently, primary and secondary screening were conducted in the same manner as in the first method. We screened more than 20,000 mutagenized F2 strains.

### Whole genome sequence

Five 6-cm NGM plates containing many adult worms were prepared. After collecting and washing the nematodes with M9 buffer, genomic DNA was purified using the GenEluteTM Mammalian Genomic DNA Miniprep Kit (Sigma, G1N10). Whole genome sequencing (WGS) was performed by BGI JAPAN. The WGS data were mapped based on the method described by Rödelsperger et al, 2020 [77] to identify mutation sites. Briefly, Illumina read data were aligned to the El Paco genome assembly [25,73] using the BWA Mem program [78]. The initial variant call was generated using the mpileup command in BCFtools [79].

### Genetic mapping

Recombinant lines used for genetic mapping were obtained by crossing light-unresponsive mutants (derived from PS312) with a male wild-type strain RSA076. Light-unresponsive F2 individuals were isolated using a light avoidance assay. After laying eggs, F2 individuals were lysed using a worm lysis buffer. To confirm the light insensitivity of the F2 individuals, a light avoidance assay was repeated on the F3 individuals. Primers were designed around marker sequences from Pristionchus.org (http://pristionchus.org), which contained insertions or deletions in PS312 and RSA076. PCR with Dream Taq Green (Thermo Fisher Scientific, K1081) was used to determine the genotype. The primers used for chromosome mapping are listed (S5 Table). Some primers were adapted from a previous study [80].

### CRISPR/Cas9 mutagenesis

To generate CRISPR knockout mutants, we followed previously described co-injection marker methods [81–83]. The CRISPR target sequences were designed using CHOP-CHOP v3 (http://chopchop.cbu.uib.no/) [84]. All tracrRNAs, crRNAs, and Cas9 proteins were synthesized by Integrated DNA Technologies (IDT). We mixed 0.5 µl of the Cas9 protein (10 µg/µl), 0.95 µl of crRNA (100 µM), and 0.9 µl of tracrRNA (100 µM), and incubated the mixture at 37 °C for 15 minutes. Using the co-CRISPR system, we combined each RNP complex containing the gRNA of *Ppa-prl-1* and a target gene. For the fluorescence marker method, we added *Ppa-egl- 20p::turboRFP* or *Ppa-eft-3p::turboRFP* (50 ng/µl) to the RNP complex and diluted with nuclease-free water up to 20 µl. The injection mixtures were microinjected into the gonads of young adult worms. The injected worms (P0) were placed individually on NGM plates. Approximately 24–48 hours later, P0 worms were removed from the plate. After 3–4 days, the F1 worms were screened for the presence of a roller phenotype or fluorescent worms. For mutation screening, a heteroduplex mobility assay was performed using microchip electrophoresis on MultiNA (Shimazu, MCE-202) or the DNA gel separation improvement agent Loupe 4 K/20 (GelBio). The genotype was determined using sanger sequencing. The identified mutants were subsequently backcrossed with the original wild-type strain (PS312) for at least three generations to eliminate off-target effects. The target sequences of the gRNA and primers are listed in S5 Table.

### Generating transgenic lines

The promoter regions were amplified using KOD One PCR Master Mix (TOYOBO, KMM-101). The lengths of the promoter sequences were as follows: *Ppa-daf-11*:794 bp; *Ppa-tax-2*:2401 bp; *Ppa-tax-4*:3001 bp*; Ppa-grk-2*:3001 bp. The promoter for *Ppa-daf-11* was constructed as described in a previous study [59]. For *Ppa-tax-2*, we used a region predicted to be a promoter in a previous study [85]. For *Ppa-tax-4* and *Ppa-grk-2*, we obtained a sequence of 3001 bp sequence upstream from the start codon. These promotors were cloned into vector containing codon optimized GFP, TurboRFP or *tetanus toxin* and *Ppa-rpl-23* 3’ UTR [82]. The plasmids and genomic DNA of PS312 were digested using *Hind*Ⅲ (pMO56, pMO59, and pMO81) or *Pst*Ⅰ (pMO74). These transgenes (3–5 ng/µl), *Ppa-egl-20p::RFP* or *Ppa-egl-20p::GFP* (50 ng/µl) as co-injection markers, and genomic DNA (60 ng/µl) were injected into the gonad of young adult worms. The transgenic animals were screened under a fluorescence microscope (M165 FC; Leica).

To identify the cell type of the amphid neurons, we followed previously described staining methods using the lipophilic dye DiI (Thermo Fisher Scientific, D3911) or Fast DiO (Thermo Fisher Scientific, D3898) [27,37]. Well-fed J2 or J3 larvae were collected in M9 buffer and centrifuged at 1500 × g for 2 min. After discarding the supernatant, worms were incubated with 150 µl of M9 containing a 1:150 dilution of FastDiO or a 1:74 dilution of DiI for 1.5 h at 20 °. The nematodes were washed twice with 1 ml of M9 buffer and crawl freely on *E. coli* seeded NGM plates for more than 1 h. Worms were immobilized on 2% agarose pads containing 5 mM levamisole or 0.3% sodium azide and covered with a cover slip. Z-stack images were obtained using a confocal microscope (Zeiss LSM900). The cell type of the amphid neurons was identified by analyzing the positional relationship between the stained cells and cells expressing the fluorescent protein. At least ten animals were observed for each reporter strain. If the fluorescence was observed in the neurons of more than 80% of the individuals, the cells were considered to express the gene.

### Statistical analysis

The Prism software package GraphPad Software 9 was used for statistical analyses. Information about the statistical tests, p-values, and n numbers is provided in the respective Figures and Figure legends. All error bars show SEM.

### Material and Data Availability

All materials generated in this study, including plasmids and worm strains, are available upon request. Any additional information required to reanalyze the data reported in this paper is available from the corresponding author upon request. The data underlying this article are available in DDBJ at https://www.ddbj.nig.ac.jp/index-e.html, and can be accessed with PRJDB 18138.

## Acknowledgments

We thank Ms. Masako Shigemori and Ms. Satoko Okazaki (Hiroshima University) for their technical support. We thank Dr. Kozue Hamao (Hiroshima University), Dr. Toshiaki Kozuka (Kanazawa University), and Dr. Takuma Sugi (Hiroshima University) for their advice regarding this study. Nematode strains and plasmids were obtained from the Caenorhabditis Genetics Center and Dr. Ralf J. Sommer (Max Planck Institute for Biology, Tübingen, Germany). This study was conducted at the Natural Science Center for Basic Research and Development at Hiroshima University. We also thank all the members of the Chihara laboratory for their kind support and Editage (www.editage.jp) for English language editing.

## Supporting information

S1 Fig. *Ppa-grk-2* is expressed in various tissues.

Representative fluorescence images of *Ppa-grk-2p::GFP*. (A) Merged image of differential interference contrast (DIC) and fluorescence, and (B) fluorescence image. GFP was expressed in pharyngeal muscles, head neurons, body wall muscles, ventral nerve cord, and tail neurons. Scale bars = 20 μm.

S2 Fig. Cilia functions are not required for light avoidance.

Blue light avoidance assay for cilia-related mutants. The mutants exhibited a normal percentage of light avoidance. One-way ANOVA Dunnett’s multiple comparison tests, compared with wild type. n.s. = not significant.

S3 Fig. Gene structures and CRISPR/Cas9 knock-out sites.

Green boxes and orange arrowheads represent exons and target regions of the gRNA, respectively. Gene structures were based on El_Paco_annotation_V3 [73]. The illustrations were created by TBtools [86].

**S1 Table. Orthogroup prediction data from OrthoFinder.**

**S2 Table. PSI-/DELTA-BLAST and tBLASTn output results.**

**S3 Table. List of *gur* genes in *P. pacificus* genome.**

**S4 Table. All strains used in this study.**

**S5 Table. All target sites and primers used in this study.**

**S1 Movie. Blue light avoidance in wild type *P. pacificus***.

The wild-type strain PS312 moved to the bottom. After 5 s, blue light (470 nm) was applied to the head. At 8 s, the worm stopped moving and initiated backward movement.

**S1 Data. LITE-1 family protein sequences.**

## Reference

1. McCulloch KJ, Macias-Muñoz A, Briscoe AD. Insect opsins and evo-devo: what have we learned in 25 years? Philos Trans R Soc Lond B Biol Sci. 2022;377: 20210288. doi:10.1098/rstb.2021.0288

2. Hagen JFD, Roberts NS, Johnston RJ Jr. The evolutionary history and spectral tuning of vertebrate visual opsins. Dev Biol. 2023;493: 40–66. doi:10.1016/j.ydbio.2022.10.014

3. Oota M, Gotoh E, Endo M, Ishida T, Matsushita T, Sawa S. Negative phototaxis in M. incognita. Int J Biol. 2017;9: 51. doi:10.5539/ijb.v9n3p51

4. Hayasaki M. Infrared light photobiostimulation mediates periodicity in Dirofilaria immitis microfilariae. J Vet Med Sci. 2020;82: 237–246. doi:10.1292/jvms.19-0156

5. Burr AH. Analysis of phototaxis in nemato des using directional statistics. J Comp Physiol. 1979;134: 85–93. doi:10.1007/BF00610280

6. Burr AH, Wagar D, Sidhu P. Ocellar pigmentation and phototaxis in the nematode Mermis nigrescens: changes during development. J Exp Biol. 2000;203: 1341–1350. doi:10.1242/jeb.203.8.1341

7. Ward A, Liu J, Feng Z, Xu XZS. Light-sensitive neurons and channels mediate phototaxis in C. elegans. Nat Neurosci. 2008;11: 916–922. doi:10.1038/nn.2155

8. Ozawa K, Shinkai Y, Kako K, Fukamizu A, Doi M. The molecular and neural regulation of ultraviolet light phototaxis and its food-associated learning behavioral plasticity in C. elegans. Neurosci Lett. 2021;770: 136384. doi:10.1016/j.neulet.2021.136384

9. Bhatla N, Horvitz HR. Light and Hydrogen Peroxide Inhibit C.elegans Feeding through Gustatory Receptor Orthologs and Pharyngeal Neurons. Neuron. 2015;85: 804–818. doi:10.1016/j.neuron.2014.12.061

10. Bhatla N, Droste R, Sando SR, Huang A, Horvitz HR. Distinct neural circuits control rhythm inhibition and spitting by the myogenic pharynx of C. elegans. Curr Biol. 2015;25: 2075–2089. doi:10.1016/j.cub.2015.06.052

11. Sando SR, Bhatla N, Lee ELQ, Robert Horvitz H. An hourglass circuit motif transforms a motor program via subcellularly localized muscle calcium signaling and contraction. Elife. 2021;10: 1–38. doi:10.7554/ELIFE.59341

12. Dipon Ghosh D, Lee D, Jin X, Robert Horvitz H, Nitabach MN. C. elegans discriminates colors to guide foraging. Science. 2021;371: 1059–1063.

13. Edwards SL, Charlie NK, Milfort MC, Brown BS, Gravlin CN, Knecht JE, et al. A novel molecular solution for ultraviolet light detection in Caenorhabditis elegans. PLoS Biol. 2008;6: 1715–1729. doi:10.1371/journal.pbio.0060198

14. Liu J, Ward A, Gao J, Dong Y, Nishio N, Inada H, et al. C. elegans phototransduction requires a G protein-dependent cGMP pathway and a taste receptor homolog. Nat Neurosci. 2010;13: 715–722. doi:10.1038/nn.2540

15. Hanson SM, Scholüke J, Liewald J, Sharma R, Ruse C, Engel M, et al. Structure-function analysis suggests that the photoreceptor LITE-1 is a light-activated ion channel. Curr Biol. 2023. doi:10.1016/j.cub.2023.07.008

16. Benton R, Himmel NJ. Structural screens identify candidate human homologs of insect chemoreceptors and cryptic Drosophila gustatory receptor-like proteins. Elife. 2023;12: e85537. doi:10.7554/eLife.85537

17. Benton R, Sachse S, Michnick SW, Vosshall LB. Atypical membrane topology and heteromeric function of Drosophila odorant receptors in vivo. PLoS Biol. 2006;4: e20. doi:10.1371/journal.pbio.0040020

18. Gong J, Yuan Y, Ward A, Feng Z, Liu J, Shawn XZ, et al. The C. elegans Taste Receptor Homolog LITE-1 Is a Photoreceptor. Cell. 2016;167: 1252–1263.e10. doi:10.1016/j.cell.2016.10.053

19. Benton R, Dessimoz C, Moi D. A putative origin of the insect chemosensory receptor superfamily in the last common eukaryotic ancestor. Elife. 2020;9: e62507. doi:10.7554/eLife.62507

20. Himmel NJ, Moi D, Benton R. Remote homolog detection places insect chemoreceptors in a cryptic protein superfamily spanning the tree of life. Curr Biol. 2023;33: 5023–5033.e4. doi:10.1016/j.cub.2023.10.008

21. Sommer RJ, Carta LK, Kim S-Y, Sternberg PWS. Morphological, genetic and molecular description of Pristionchus pacificus sp. n. (Nematoda:Neodiplogastridae). Fundamental and applied Nematology. 1996. pp. 511–521. Available: https://core.ac.uk/download/pdf/39853320.pdf

22. Sommer RJ, Sternberg PW. Apoptosis and change of competence limit the size of the vulva equivalence group in Pristionchus pacificus: a genetic analysis. Curr Biol. 1996;6: 52–59. doi:10.1016/S0960-9822(02)00421-9

23. Schroeder NE. Introduction to Pristionchus pacificus anatomy. J Nematol. 2021;53. doi:10.21307/jofnem-2021-091

24. Dieterich C, Clifton SW, Schuster LN, Chinwalla A, Delehaunty K, Dinkelacker I, et al. The Pristionchus pacificus genome provides a unique perspective on nematode lifestyle and parasitism. Nat Genet. 2008;40: 1193–1198. doi:10.1038/ng.227

25. Rödelsperger C, Meyer JM, Prabh N, Lanz C, Bemm F, Sommer RJ. Single-Molecule Sequencing Reveals the Chromosome-Scale Genomic Architecture of the Nematode Model Organism Pristionchus pacificus. Cell Rep. 2017;21: 834–844. doi:10.1016/j.celrep.2017.09.077

26. Witte H, Moreno E, Rödelsperger C, Kim J, Kim J-S, Streit A, et al. Gene inactivation using the CRISPR/Cas9 system in the nematode Pristionchus pacificus. Dev Genes Evol. 2015;225: 55–62. doi:10.1007/s00427-014-0486-8

27. Hong RL, Riebesell M, Bumbarger DJ, Cook SJ, Carstensen HR, Sarpolaki T, et al. Evolution of neuronal anatomy and circuitry in two highly divergent nematode species. Elife. 2019;8: 1–34. doi:10.7554/eLife.47155

28. Bumbarger DJ, Riebesell M, Rödelsperger C, Sommer RJ. System-wide rewiring underlies behavioral differences in predatory and bacterial-feeding nematodes. Cell. 2013;152: 109–119. doi:10.1016/j.cell.2012.12.013

29. Okumura M, Wilecki M, Sommer RJ. Serotonin Drives Predatory Feeding Behavior via Synchronous Feeding Rhythms in the Nematode Pristionchus pacificus. G3. 2017;7: 3745–3755. doi:10.1534/g3.117.300263

30. Lightfoot JW, Wilecki M, Rödelsperger C, Moreno E, Susoy V, Witte H, et al. Small peptide-mediated self-recognition prevents cannibalism in predatory nematodes. Science. 2019;364: 86–89. doi:10.1126/science.aav9856

31. Kroetz SM, Srinivasan J, Yaghoobian J, Sternberg PW, Hong RL. The cGMP signaling pathway affects feeding behavior in the necromenic nematode Pristionchus pacificus. PLoS One. 2012;7: e34464. doi:10.1371/journal.pone.0034464

32. Hong RL, Witte H, Sommer RJ. Natural variation in Pristionchus pacificus insect pheromone attraction involves the protein kinase EGL-4. Proc Natl Acad Sci U S A. 2008;105: 7779–7784. doi:10.1073/pnas.0708406105

33. Hiramatsu F, Lightfoot JW. Kin-recognition and predation shape collective behaviors in the cannibalistic nematode Pristionchus pacificus. PLoS Genet. 2023;19: e1011056. doi:10.1371/journal.pgen.1011056

34. Hong RL, Sommer RJ. Chemoattraction in Pristionchus nematodes and implications for insect recognition. Curr Biol. 2006;16: 2359–2365. doi:10.1016/j.cub.2006.10.031

35. Ishita Y, Chihara T, Okumura M. Different combinations of serotonin receptors regulate predatory and bacterial feeding behaviors in the nematode Pristionchus pacificus. G3. 2021;11. doi:10.1093/g3journal/jkab011

36. Lo W-S, Roca M, Dardiry M, Mackie M, Eberhardt G, Witte H, et al. Evolution and Diversity of TGF-β Pathways are Linked with Novel Developmental and Behavioral Traits. Mol Biol Evol. 2022;39. doi:10.1093/molbev/msac252

37. Quach KT, Chalasani SH. Flexible reprogramming of Pristionchus pacificus motivation for attacking Caenorhabditis elegans in predator-prey competition. Curr Biol. 2022; 1–14. doi:10.1016/j.cub.2022.02.033

38. Moreno E, Sieriebriennikov B, Witte H, Rödelsperger C, Lightfoot JW, Sommer RJ. Regulation of hyperoxia-induced social behaviour in Pristionchus pacificus nematodes requires a novel cilia-mediated environmental input. Sci Rep. 2017;7. doi:10.1038/s41598-017-18019-0

39. Pratx L, Rancurel C, Da Rocha M, Danchin EGJ, Castagnone-Sereno P, Abad P, et al. Genome-wide expert annotation of the epigenetic machinery of the plant-parasitic nematodes Meloidogyne spp., with a focus on the asexually reproducing species. BMC Genomics. 2018;19: 321. doi:10.1186/s12864-018-4686-x

40. Brown AL, Meiborg AB, Franz-Wachtel M, Macek B, Gordon S, Rog O, et al. Characterization of the Pristionchus pacificus “epigenetic toolkit” reveals the evolutionary loss of the histone methyltransferase complex PRC2. Genetics. 2024; iyae041. doi:10.1093/genetics/iyae041

41. Yau K-W, Hardie RC. Phototransduction motifs and variations. Cell. 2009;139: 246–264. doi:10.1016/j.cell.2009.09.029

42. Gühmann M, Porter ML, Bok MJ. The Gluopsins: Opsins without the Retinal Binding Lysine. Cells. 2022;11. doi:10.3390/cells11152441

43. Deppisch P, Helfrich-Förster C, Senthilan PR. The Gain and Loss of Cryptochrome/Photolyase Family Members during Evolution. Genes. 2022;13. doi:10.3390/genes13091613

44. Howe KL, Bolt BJ, Shafie M, Kersey P, Berriman M. WormBase ParaSite - a comprehensive resource for helminth genomics. Mol Biochem Parasitol. 2017;215: 2–10. doi:10.1016/j.molbiopara.2016.11.005

45. Xiang Y, Yuan Q, Vogt N, Looger LL, Jan LY, Jan YN. Light-avoidance-mediating photoreceptors tile the Drosophila larval body wall. Nature. 2010;468: 921–926. doi:10.1038/nature09576

46. Schäffer AA, Aravind L, Madden TL, Shavirin S, Spouge JL, Wolf YI, et al. Improving the accuracy of PSI-BLAST protein database searches with composition-based statistics and other refinements. Nucleic Acids Res. 2001;29: 2994–3005. doi:10.1093/nar/29.14.2994

47. Altschul SF, Madden TL, Schäffer AA, Zhang J, Zhang Z, Miller W, et al. Gapped BLAST and PSI-BLAST: a new generation of protein database search programs. Nucleic Acids Res. 1997;25: 3389–3402. doi:10.1093/nar/25.17.3389

48. Boratyn GM, Schäffer AA, Agarwala R, Altschul SF, Lipman DJ, Madden TL. Domain enhanced lookup time accelerated BLAST. Biol Direct. 2012;7: 12. doi:10.1186/1745-6150-7-12

49. Wang JK, McDowell JH, Hargrave PA. Site of attachment of 11-cis-retinal in bovine rhodopsin. Biochemistry. 1980;19: 5111–5117. doi:10.1021/bi00563a027

50. Troemel ER, Chou JH, Dwyer ND, Colbert HA, Bargmann CI. Divergent seven transmembrane receptors are candidate chemosensory receptors in C. elegans. Cell. 1995;83: 207–218. doi:10.1016/0092-8674(95)90162-0

51. Gurevich VV, Gurevich EV. GPCR Signaling Regulation: The Role of GRKs and Arrestins. Front Pharmacol. 2019;10: 125. doi:10.3389/fphar.2019.00125

52. Kühn H, Wilden U. Deactivation of photoactivated rhodopsin by rhodopsin-kinase and arrestin. J Recept Res. 1987;7: 283–298. doi:10.3109/10799898709054990

53. Zhao X, Huang J, Khani SC, Palczewski K. Molecular forms of human rhodopsin kinase (GRK1). J Biol Chem. 1998;273: 5124–5131. doi:10.1074/jbc.273.9.5124

54. Chen CK, Zhang K, Church-Kopish J, Huang W, Zhang H, Chen YJ, et al. Characterization of human GRK7 as a potential cone opsin kinase. Mol Vis. 2001;7: 305–313. Available: https://www.ncbi.nlm.nih.gov/pubmed/11754336

55. Maeda T, Imanishi Y, Palczewski K. Rhodopsin phosphorylation: 30 years later. Prog Retin Eye Res. 2003;22: 417–434. doi:10.1016/s1350-9462(03)00017-x

56. Tachibanaki S, Arinobu D, Shimauchi-Matsukawa Y, Tsushima S, Kawamura S. Highly effective phosphorylation by G protein-coupled receptor kinase 7 of light-activated visual pigment in cones. Proc Natl Acad Sci U S A. 2005;102: 9329–9334. doi:10.1073/pnas.0501875102

57. Fukuto HS, Ferkey DM, Apicella AJ, Lans H, Sharmeen T, Chen W, et al. G Protein- Coupled Receptor Kinase Function Is Essential for Chemosensation in C. elegans. Neuron. 2004;42: 581–593.

58. Wood JF, Wang J, Benovic JL, Ferkey DM. Structural domains required for Caenorhabditis elegans G protein-coupled receptor kinase 2 (GRK-2) function in Vivo. J Biol Chem. 2012;287: 12634–12644. doi:10.1074/jbc.M111.336818

59. Lenuzzi M, Witte H, Riebesell M, Rödelsperger C, Hong RL, Sommer RJ. Influence of environmental temperature on mouth-form plasticity in Pristionchus pacificus acts through daf-11-dependent cGMP signaling. J Exp Zool B Mol Dev Evol. 2021; 1–11. doi:10.1002/jez.b.23094

60. Moreno E, Lenuzzi M, Rödelsperger C, Prabh N, Witte H, Roeseler W, et al. DAF- 19/RFX controls ciliogenesis and influences oxygen-induced social behaviors in Pristionchus pacificus. Evol Dev. 2018;20: 233–243. doi:10.1111/ede.12271

61. Moreno E, Lightfoot JW, Lenuzzi M, Sommer RJ. Cilia drive developmental plasticity and are essential for efficient prey detection in predatory nematodes. Proc Biol Sci. 2019;286: 20191089. doi:10.1098/rspb.2019.1089

62. Schiavo G, Benfenati F, Poulain B, Rossetto O, Polverino de Laureto P, DasGupta BR, et al. Tetanus and botulinum-B neurotoxins block neurotransmitter release by proteolytic cleavage of synaptobrevin. Nature. 1992;359: 832–835. doi:10.1038/359832a0

63. Jin Y, Jorgensen E, Hartwieg E, Horvitz HR. The Caenorhabditis elegans gene unc-25 encodes glutamic acid decarboxylase and is required for synaptic transmission but not synaptic development. J Neurosci. 1999;19: 539–548. doi:10.1523/JNEUROSCI.19-02-00539.1999

64. Serrano-Saiz E, Poole RJ, Felton T, Zhang F, De La Cruz ED, Hobert O. Modular control of glutamatergic neuronal identity in C. elegans by distinct homeodomain proteins. Cell. 2013;155: 659–673. doi:10.1016/j.cell.2013.09.052

65. Boccuni I, Fairless R. Retinal Glutamate Neurotransmission: From Physiology to Pathophysiological Mechanisms of Retinal Ganglion Cell Degeneration. Life. 2022;12. doi:10.3390/life12050638

66. Brenner S. The genetics of Caenorhabditis elegans. Genetics. 1974;77: 71–94. doi:10.1093/genetics/77.1.71

67. Davis P, Zarowiecki M, Arnaboldi V, Becerra A, Cain S, Chan J, et al. WormBase in 2022-data, processes, and tools for analyzing Caenorhabditis elegans. Genetics. 2022;220. doi:10.1093/genetics/iyac003

68. Sternberg PW, Van Auken K, Wang Q, Wright A, Yook K, Zarowiecki M, et al. WormBase 2024: status and transitioning to Alliance infrastructure. Genetics. 2024;227. doi:10.1093/genetics/iyae050

69. Öztürk-Çolak A, Marygold SJ, Antonazzo G, Attrill H, Goutte-Gattat D, Jenkins VK, et al. FlyBase: updates to the Drosophila genes and genomes database. Genetics. 2024;227. doi:10.1093/genetics/iyad211

70. Mistry J, Chuguransky S, Williams L, Qureshi M, Salazar GA, Sonnhammer ELL, et al. Pfam: The protein families database in 2021. Nucleic Acids Res. 2021;49: D412–D419. doi:10.1093/nar/gkaa913

71. Paysan-Lafosse T, Blum M, Chuguransky S, Grego T, Pinto BL, Salazar GA, et al. InterPro in 2022. Nucleic Acids Res. 2023;51: D418–D427. doi:10.1093/nar/gkac993

72. Emms DM, Kelly S. OrthoFinder: phylogenetic orthology inference for comparative genomics. Genome Biol. 2019;20: 238. doi:10.1186/s13059-019-1832-y

73. Athanasouli M, Witte H, Weiler C, Loschko T, Eberhardt G, Sommer RJ, et al. Comparative genomics and community curation further improve gene annotations in the nematode Pristionchus pacificus. BMC Genomics. 2020;21: 1–9. doi:10.1186/s12864-020-07100-0

74. Katoh K, Standley DM. MAFFT multiple sequence alignment software version 7: improvements in performance and usability. Mol Biol Evol. 2013;30: 772–780. doi:10.1093/molbev/mst010

75. Zhou L, Feng T, Xu S, Gao F, Lam TT, Wang Q, et al. ggmsa: a visual exploration tool for multiple sequence alignment and associated data. Brief Bioinform. 2022;23. doi:10.1093/bib/bbac222

76. Silva AP da. Pristionchus pacificus genetic protocols. WormBook. 2005; 1–8. doi:10.1895/wormbook.1.114.1

77. Rödelsperger C. A simplified workflow for the analysis of whole-genome sequencing data from Pristionchus pacificus mutant lines. bioRxiv. 2020. p. 2020.11.12.379388. doi:10.1101/2020.11.12.379388

78. Li H, Durbin R. Fast and accurate short read alignment with Burrows-Wheeler transform. Bioinformatics. 2009;25: 1754–1760. doi:10.1093/bioinformatics/btp324

79. Li H. A statistical framework for SNP calling, mutation discovery, association mapping and population genetical parameter estimation from sequencing data. Bioinformatics. 2011;27: 2987–2993. doi:10.1093/bioinformatics/btr509

80. Ishita Y, Onodera A, Ekino T, Chihara T, Okumura M. Co-option of an Astacin Metalloprotease Is Associated with an Evolutionarily Novel Feeding Morphology in a Predatory Nematode. Mol Biol Evol. 2023;40. doi:10.1093/molbev/msad266

81. Nakayama KI, Ishita Y, Chihara T, Okumura M. Screening for CRISPR/Cas9-induced mutations using a co-injection marker in the nematode Pristionchus pacificus. Dev Genes Evol. 2020;230: 257–264. doi:10.1007/s00427-020-00651-y

82. Han Z, Lo W-S, Lightfoot JW, Witte H, Sun S, Sommer RJ. Improving Transgenesis Efficiency and CRISPR-Associated Tools Through Codon Optimization and Native Intron Addition in Pristionchus Nematodes. Genetics. 2020; genetics.303785.2020. doi:10.1534/genetics.120.303785

83. Hiraga H, Ishita Y, Chihara T, Okumura M. Efficient visual screening of CRISPR/Cas9 genome editing in the nematode Pristionchus pacificus. Dev Growth Differ. 2021;63: 488–500. doi:10.1111/dgd.12761

84. Labun K, Montague TG, Krause M, Torres Cleuren YN, Tjeldnes H, Valen E. CHOPCHOP v3: expanding the CRISPR web toolbox beyond genome editing. Nucleic Acids Res. 2019;47: W171–W174. doi:10.1093/nar/gkz365

85. Werner MS, Sieriebriennikov B, Prabh N, Loschko T, Lanz C, Sommer RJ. Young genes have distinct gene structure, epigenetic profiles, and transcriptional regulation. Genome Res. 2018;28: 1675–1687. doi:10.1101/gr.234872.118

86. Chen C, Chen H, Zhang Y, Thomas HR, Frank MH, He Y, et al. TBtools: An Integrative Toolkit Developed for Interactive Analyses of Big Biological Data. Mol Plant. 2020;13: 1194–1202. doi:10.1016/j.molp.2020.06.009

